# SILAC-Site: A streamlined workflow for determining phosphopeptide stoichiometries

**DOI:** 10.64898/2026.05.28.728441

**Authors:** Katharina D. Faisst, Ludwig R. Sinn, Kate Lau, Lukasz Szyrwiel, Juri Rappsilber, Vadim Demichev

## Abstract

Mass spectrometry-based proteomics enables in-depth investigation of protein phosphorylation, quantifying tens of thousands of phosphosites per sample following phosphopeptide enrichment. However, critical information on phosphosite occupancy (stoichiometry) is typically lost during the phospho-enrichment process. Here, we introduce SILAC-Site, a SILAC-based fractionation-free and chemical labelling-free workflow for direct phosphosite stoichiometry evaluation using stable isotope labeling and phosphatase treatment. By acquiring treated or untreated peptides together with their heavy-labelled dephosphorylated counterparts within the same LC-MS runs, this approach provides internally controlled stoichiometry estimates compatible with high-throughput data-independent acquisition proteomics. Applying SILAC-Site to *S. cerevisiae*, we show that the majority of phosphopeptides identified only after enrichment possess low stoichiometries, and that inferred stoichiometry strongly correlates with the direct detection of phosphopeptides in samples without enrichment. Based on these findings, we propose the analysis of samples without enrichment as a simple complementary addition to a typical phosphoproteomics workflow, facilitating recovery of phosphorylation stoichiometry information.

## Introduction

Data-independent acquisition (DIA) proteomics has gained prominence through advances in mass spectrometry (MS) instrumentation and computational data processing and is particularly effective for quantitative analysis of post-translational modifications such as protein phosphorylation.^1–3^ While phosphopeptide enrichment methods enable effective comparisons of phosphorylation levels between conditions, information on phosphosites stoichiometries remains largely inaccessible.^4,5^ Indeed, enrichment-based phosphoproteomics cannot directly measure stoichiometries, typically relying on indirect computational estimates.^6^

To address this problem, Wu *et al* examined the effect of phosphatase treatment on the quantities of unmodified peptides, in samples without enrichment, demonstrating the feasibility of direct phosphopeptide stoichiometry measurement in yeast.^4^ In later works, this strategy was extended to human cells and adapted for isobaric labelling, to enable multiplexed analysis while maintaining the core principle of comparing phosphatase-treated and untreated samples.^7,8^ However, these methods relied on offline fractionation and data-dependent acquisition (DDA), making the workflows less reproducible, labor-intensive and requiring large input sample amounts. Furthermore, chemical labelling strategies such as dimethyl labelling, mTRAQ or TMT, can introduce systematic biases through incomplete labelling or peptide-specific variability.^9–12^

Here, we leverage recent advances in DIA proteomics to simplify the workflow and eliminate the need for chemical labelling. We introduce SILAC-Site, a method for quantifying protein and peptide dephosphorylation to estimate phosphosite stoichiometry using stable isotope labelling by amino acids in cell culture (SILAC). SILAC ensures complete and consistent isotope incorporation across the proteome.^13,14^ This enables accurate quantification based on the ratios between light and heavy isotopologues, reducing the impact of LC-MS-related variation, such as matrix and ionisation effects. Integrated with our recently introduced Slice-PASEF acquisition technology,^15^ aimed at maximising quantitative precision and data completeness, SILAC-Site achieves high reproducibility while retaining robustness against LC-MS-derived errors.

Applied to yeast (*S. cerevisiae*), SILAC-Site revealed that phosphopeptides directly detectable by DIA without enrichment tend to exhibit higher stoichiometries, whereas phosphopeptides detected only following enrichment predominantly represent low-stoichiometry phosphorylation sites. Our results support the analysis of non-enriched samples as a simple complement to phosphopeptide enrichment workflows for recovering phosphorylation stoichiometry information.

## Results

### Design of the SILAC-Site workflow

Phosphatase-based phosphorylation-site stoichiometry analysis relies on quantifying changes in peptide abundance before and after enzymatic dephosphorylation. Here, an increase in the quantity of an unmodified peptide species upon dephosphorylation reflects the cumulative occupancy of phosphorylation sites on the peptide. In SILAC-Site, dephosphorylated heavy-isotope labelled peptides are first mixed with light isotope peptides that retain their native phosphorylation state, then split and either treated with calf intestinal alkaline phosphatase (CIP) or left untreated (Figure 1a, left panel).

**Figure 1.**
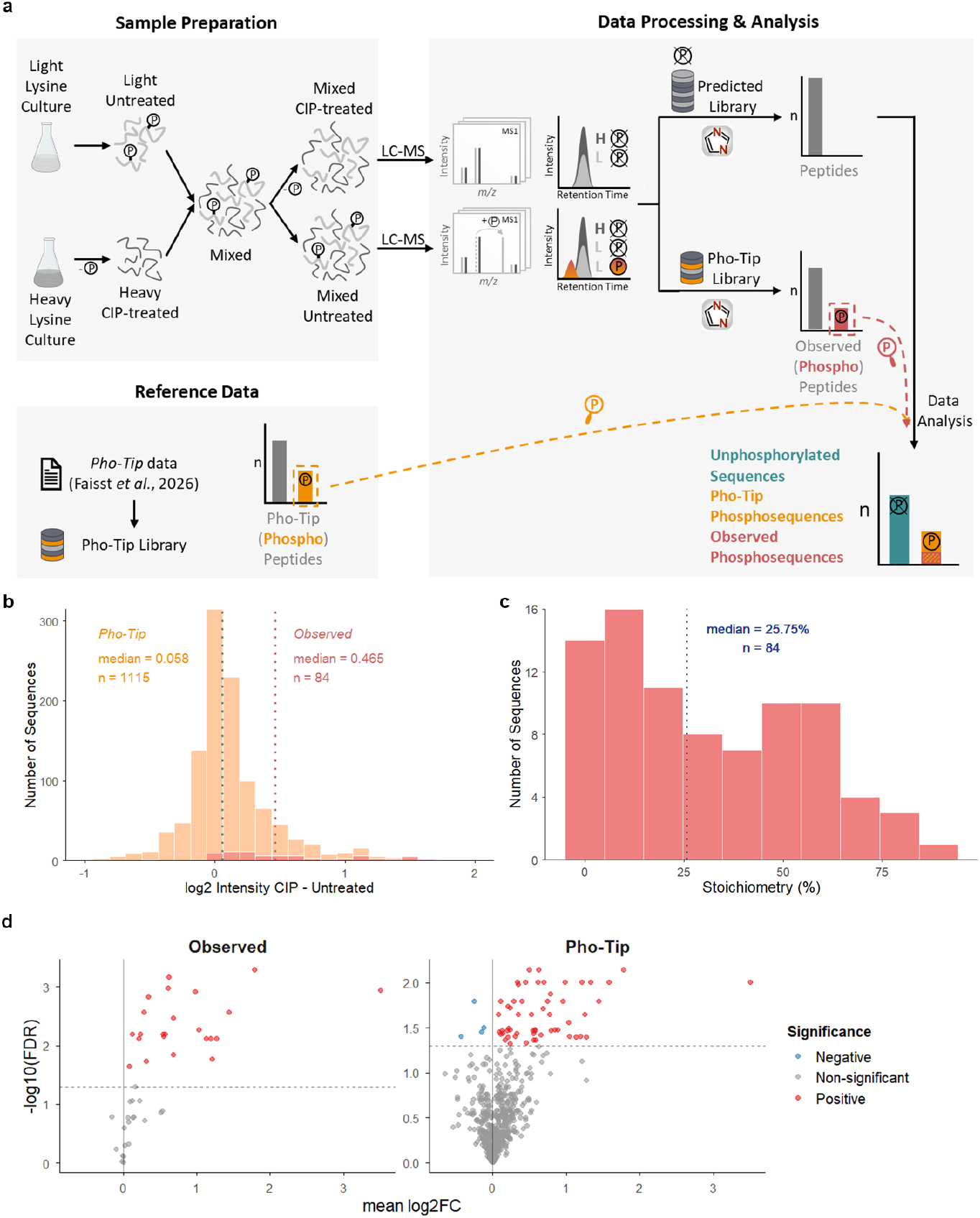
SILAC-Site workflow and data analysis. **a)** Yeast were cultured in media supplemented with light (L) or heavy (H) lysine. After digestion, samples were treated with CIP (H CIP) or left untreated (L Untreated). H CIP samples were mixed with L Untreated, split and treated with CIP (Mixed CIP) or left untreated (Mixed Untreated). Following LC-MS analysis, data were searched against a fully predicted library (no variable modifications) and against the Pho-Tip^16^ spectral library for “observed phosphopeptides” detection. Predicted library output was filtered for unmodified peptides matching the observed phosphopeptides and/or all phosphopeptides in the Pho-Tip library (“Pho-Tip phosphopeptides”, orange), yielding “observed phosphosequences” (red), “Pho-Tip phosphosequences” (orange), and “unphosphorylated sequences” (turquoise). **b)** Distribution of log-scale (L/H) intensity shifts upon CIP treatment for non-observed Pho-Tip and observed phosphosequences. **c)** Inferred phosphopeptide stoichiometry (Methods) distribution for observed phosphosequences, based on the intensity shifts. **d)** Volcano plot showing changes upon CIP treatment, across observed phosphosequences (left) and Pho-Tip phosphosequences (right). Each point represents one peptide sequence, detected in all technical replicates (n = 5). The x-axis shows the mean log-scale (L/H) intensity shift upon CIP treatment, and the y-axis shows −log10 adjusted p-values from one-sample t-tests. Phosphosequences with significant positive shifts (FDR < 0.05, log_2_FC > 0) are coloured in red, those with negative shifts (log_2_FC < 0) in blue, and non-significant in grey. The dashed line marks the significance threshold (-log_10_(0.05)).

Unlike pure label-free quantification, where matrix effects associated with sample preparation may cause ionisation differences between acquisitions and introduce unknown biases to the inferred peptide quantities, in SILAC-Site both, the light unmodified peptide, CIP-treated or untreated, and its CIP-treated heavy counterpart are measured simultaneously. While fluctuations in LC-MS response affect both light and heavy ion species, the light-to-heavy ratios remain a robust readout, enabling quantitative comparison of measured peptide intensities between CIP-treated and untreated light fractions.

Thus, the SILAC-Site workflow provides proteome-wide sequence-specific internal standards to control for technical variation, while avoiding the limitations associated with chemical labelling methods by ensuring quasi complete proteome-wide metabolic label incorporation. These features establish SILAC-Site as a robust platform for stoichiometry estimation compatible with high-throughput data-independent acquisition (DIA) workflows.

### Application of SILAC-Site to yeast phosphoproteomics

We applied the SILAC-Site workflow to *S. cerevisiae* growing to exponential phase, with data acquired on an Evosep One liquid chromatography system coupled to a timsTOF Ultra mass spectrometer operated in a 2-frame Slice-PASEF mode^15^ to ensure high sensitivity and quantitative precision (Methods).

The LC-MS data were searched against an *in silico* predicted spectral library without variable modifications, to quantify unmodified peptides and determine their light-to-heavy (L/H) ratios (Figure 1a). In parallel, the data were also searched using the “Pho-Tip” yeast phosphopeptide spectral library from our previous study,^16^ to identify phosphopeptides directly detectable in the non-enriched SILAC-Site samples (“observed phosphopeptides”). The unmodified peptides were matched to these directly observed phosphopeptides (resulting in “observed phosphosequences”) or, more broadly, to all phosphopeptides annotated in the Pho-Tip library (“Pho-Tip phosphosequences”). All remaining unmodified peptides detected using the predicted library were classified as “unphosphorylated sequences”. Since phosphate groups are removed upon CIP treatment, stoichiometry estimation was performed at the level of peptide sequences rather than specific modification sites. To facilitate interpretation of the data, we used consistent terminology and colour scheme throughout the manuscript when referring to the above classes of peptides (Figure S1).

First, we confirmed robust LC-MS performance. Indeed, we observed consistent identification coverage and good quantitative precision across replicates and labelling channels (Figures S2, S3a,b). A quantitative comparison of untreated and CIP-treated samples revealed no overall shift in background peptide signals. The quantities’ distributions remained consistent across peptides without phosphorylation evidence as well as non-phosphorylatable peptides lacking Ser/Thr/Tyr (STY) residues (Figure S4a-c). Further, analysing the observed phosphopeptides, we confirmed efficient CIP-mediated dephosphorylation (Figure S5a,b).

### Phosphopeptides detectable without enrichment show high stoichiometries

Given the typically impaired chromatographic separation as well as lower ionisation efficiency of phosphopeptides in comparison to their unmodified counterparts,^16–18^ we speculated that the observed phosphopeptides may correspond to high-stoichiometry sites. Indeed, examination of observed phosphosequences in Pho-Tip data revealed high inferred occupancies, with a median value of 25.75% (Figure 1b,c, Methods). In contrast, when we analysed intensity shifts upon CIP treatment for non-observed Pho-Tip phosphosequences, most showed no change beyond the level expected from measurement variability (Figure 1b). Indeed, statistical testing of peptide sequences affected by CIP treatment showed that 53% of observed phosphosequences detected in all replicates had significant shifts to higher signals (n = 23/43), compared with only 8% of Pho-Tip phosphosequences (n = 56/663; Figure 1d).

We further noted that peptide sequences with high stoichiometries may only be detectable upon CIP treatment. Consistent with this, 22% (n = 24/109, L channel) of observed phosphosequences were identified exclusively in their dephosphorylated form after CIP treatment (Figure S5b). To take these peptides into account, we therefore performed an additional analysis where missing intensity values in the light channel of both CIP-treated and untreated samples were imputed using half-minimum value imputation to approximate signals below the detection limit (Figure S6). Imputation minimally affected Pho-Tip phosphosequences but increased the median inferred stoichiometry for observed phosphosequences to 42.4 % (Figure S6a,b).

We next examined the stoichiometries inferred using exclusively the light channel data, that is, without leveraging the internal heavy-labelled controls provided by SILAC-Site. This yielded a similar stoichiometry distribution and showed strong overall correlation with the SILAC-Site measurements (Figure S6c,d), although individual data points obtained using only light channel data are arguably less reliable.

We further performed an orthogonal validation of SILAC-Site. For this, we compared the stoichiometries inferred by SILAC-Site for observed phosphosequences to stoichiometry estimates obtained based on the intensities of phosphorylated and unmodified forms of the same peptides detected in the SILAC-Site experiment (Figure 1c). Here, we further leveraged the relative ionisation efficiencies of these peptides that we have measured previously.^16^ This comparison showed a strong agreement between the two stoichiometry estimates (Figure S7).

Finally, we compared our results with those of Wu *et al*.^*4*^ We observed that SILAC-Site sequences overlapping with phosphosequences detected by Wu *et al* showed no definitive intensity shifts upon CIP treatment (Figure S8a), while the two datasets exhibited limited overlap in identified phosphosequences and only moderate correlation of CIP treatment effects among shared observed phosphosequences (Figure S8b,c). These differences may reflect differences in acquisition strategies (DIA vs. DDA) and experimental design, including labelling approaches (SILAC vs. dimethyl labelling) and growth conditions (synthetic minimal vs. yeast extract peptone dextrose).

To evaluate the inter-replicate variability, we examined the log-space standard deviation distributions for observed phosphosequences, Pho-Tip phosphosequences as well as phosphosequences considered by Wu *et al*,^*4*^ including analyses with and without precursor filtering by a QuantUMS^19^ score reported by DIA-NN that reflects quantitative reliability, as well as with or without imputation (Figure S8d-g). Overall, these analyses support the robustness of SILAC-Site stoichiometry estimates while highlighting the increased quantitative precision achievable in deeply fractionated workflows.

## Discussion

In SILAC-Site, DIA proteomics combined with stable isotope labelling enables quantification of phosphorylation stoichiometry at the peptide level. In particular, the SILAC internal standards facilitate stoichiometry calculations regardless of the presence of undesired technical signal variation. In combination with Slice-PASEF ion mobility-resolved mass spectrometry, this approach achieves high quantitative precision and data completeness without requiring chemical labelling of peptides or offline sample fractionation. Thus, SILAC-site can be readily integrated into conventional phosphoproteomics workflows.

We note that the use of heavy-labelled internal controls here is conceptually similar to the Super-SILAC strategy,^20,21^ wherein labelled standards serve as an internal quantitative reference. Likewise, unlike targeted phosphopeptide standard mixes that are restricted to predefined peptides, SILAC-Site enables signal normalisation across the entire proteome.

Typically, only tens to hundreds of phosphorylated peptides are identified without enrichment using single-shot DIA, in contrast to more than one hundred thousand unmodified peptides detectable in human cell lysates using state of the art workflows.^22^ Given that phosphopeptide ionisation efficiencies are, on average, reduced less than two-fold relative to their unphosphorylated counterparts,^16^ including on modern instruments, the vast majority of phosphopeptides, and consequently phosphosites, are likely to occur at low stoichiometry.

Consistent with this view, we found that phosphopeptides directly detectable without enrichment in *S. cerevisiae* generally correspond to higher inferred stoichiometries. In fact, the number of phosphopeptides with statistically significant positive stoichiometries was comparable to the number of peptides directly detectable in phosphorylated form in the same samples. In addition, stoichiometries inferred using SILAC-Site correlated well with estimates derived from measured intensities of phosphorylated and unphosphorylated peptide counterparts after correcting for relative ionisation efficiencies.^16^ Together, these observations confirm that peptides detectable in unmodified but not phosphorylated form are unlikely to exist at high phosphorylation stoichiometry.

In our analysis, we could only examine the stoichiometries for phosphopeptides that are either directly detectable as phosphorylated or have their unmodified counterparts detectable in non-enriched samples. Accordingly, our conclusions are necessarily limited to the subset of peptides derived from sufficiently abundant proteins and with sequences that ionise sufficiently well. At the same time, phosphopeptide enrichment is potentially capable of recovering even low-abundant and poor-ionising phosphopeptides. While we are not aware of any reasons why such poorly-ionising phosphopeptides would have a substantially differing distribution of stoichiometries, this remains an important limitation of the workflow.

Comparison with the stoichiometry measurements reported by Wu *et al*^*4*^ revealed limited overlap between detected phosphopeptides as well as low correlation between inferred stoichiometries. In addition, SILAC-Site generally indicated lower stoichiometries across the analysed phosphopeptides than reported by Wu and colleagues. The reasons for this discrepancy remain unclear but may reflect differences in growth conditions, as Wu *et al* analysed yeast grown in rich medium, whereas the current study used synthetic minimal medium.

While SILAC-Site offers high precision without fractionation, its reliance on metabolic labelling limits applicability to samples compatible with SILAC.^21^ At the same time, direct phosphopeptide searches in non-enriched samples show strong agreement with SILAC-Site measurements and may therefore provide a broadly applicable alternative for direct estimation of stoichiometries.

A deeper understanding of protein phosphorylation and its regulation requires knowledge of individual site occupancies.^23^ It has been argued that high phosphosite stoichiometry is often associated with functional importance of the respective site.^24^ Studies by Olsen *et al* and Bekker-Jensen *et al* show that such sites are enriched during dynamic processes like mitosis or EGF signaling.^6,24,25^ Wu and colleagues further note that these high-occupancy sites commonly occur in acidic motifs and nuclear regions, linking stoichiometry to critical cellular functions like chromatin silencing and cytokinesis.^4^

In this work, we have demonstrated that phosphosite stoichiometry can be estimated directly from non-enriched samples, either through SILAC-Site or, in case phosphopeptide ionisation efficiencies are known, through direct detection of phosphorylated peptides. Because this analysis requires only limited additional sample material and instrument time, it can be readily incorporated into phosphoproteomics workflows and may provide valuable complementary information for interpreting phosphorylation dynamics.

## Methods

### SILAC labelling and cell harvest light & heavy

Yeast *(S. cerevisiae)* BY4741 Δlys2^26^ cells were cultured in synthetic minimal (SM) medium supplemented with either light (Lys-0) or heavy (Lys-8) lysine to generate non-labelled (Light) and heavy-labelled (Heavy) samples, respectively. Cells were streaked onto SM agar plates containing light lysine and incubated for three days at 30 °C.

SM liquid media were prepared for both light lysine (6.7 g/l yeast nitrogen base (YNB) without amino acids, 2% glucose, 30 mg/l Lys-0 lysine) and heavy lysine conditions (6.7 g/l YNB without amino acids, 2% glucose, 30 mg/l Lys-8 lysine). Separate pre-cultures were inoculated and grown overnight at 30 °C, 300 rpm. Cultures were then incubated for approximately 16 h at 30 °C, 300 rpm.

Cells were harvested at late-log phase (OD = 2) by centrifugation for 15 min at 20,000 x g at 3 °C. Pellets were washed twice with ice-cold phosphate buffered saline (PBS). Resuspended cells were centrifuged at 20,000 x g at 3 °C, supernatants were discarded. Final pellets were immediately frozen at -80 °C until further processing.

### Cell lysis light & heavy samples

Yeast cell pellets from light and heavy samples were washed twice with ice-cold Milli-Q water and centrifuged for 2 min at 20,000 x g at 4 °C. The supernatant was discarded, the pellets were resuspended in 0.2 M sodium hydroxide (NaOH) and incubated on ice for 15 min. Cells were centrifuged again for 2 min at 20,000 x g at 4 °C, and the supernatant was removed. Pellets were then resuspended in lysis buffer (7 M Urea, 0.1 M ammonium bicarbonate (ABC)) containing PhosphataseArrest™ I (G-Biosciences, St. Louis, MO, USA). The suspension was split equally into two tubes and mixed with glass beads. Cell disruption was performed using the SPEX CertiPrep™ Geno/Grinder 2 (SPEX SamplePrep, Metuchen, NJ, USA) at 3,000 rpm for 30 s, followed by two 5 min cycles at 1500 rpm, with cooling on ice between cycles. Lysates were centrifuged for 15 min at 20,000 x g at 4°C. The supernatants were collected and centrifuged again for 1 h at 20,000 x g at 4 °C. Protein concentrations were determined using the Pierce™ BCA Protein Assay Kit (Thermo Fisher Scientific, Waltham, MA, USA).

### Protein reduction, alkylation and digestion light & heavy samples

Protein disulfide bonds were reduced with 5 mM dithiothreitol (DTT) for 30 min at room temperature (RT). Samples were briefly centrifuged, vortexed, and cooled on ice for 5 min. Free cysteine residues were then alkylated with 10 mM iodoacetamide (IAA) for 20 min at RT in the dark. The reaction was quenched by adding 5 mM DTT and incubating for 10 min at RT in the dark. The cell lysate was diluted 1:5 with 0.1 M ABC. Trypsin was added at an enzyme-to-protein ratio of 1:100 (m/m), and samples were briefly vortexed and centrifuged. Digestion was performed overnight at 37 °C. The reaction was stopped by acidifying with trifluoroacetic acid (TFA) to pH 2-3. After 15 min incubation at RT, the samples were centrifuged for 15 min at 20,000 x g at 4 °C, and the supernatant was collected.

### Peptide desalting light & heavy samples

Peptide desalting was performed using solid-phase extraction (SPE) C18 cartridges (Waters Corporation, Milford, MA, USA) according to the manufacturer’s instructions. In brief, cartridges were activated with methanol (MeOH) and equilibrated with buffer B (80% acetonitrile (ACN), 0.1% TFA) followed by 3 x buffer A (0.1% TFA). Samples were loaded onto the cartridges and centrifuged at 8 x g for 3-5 min. The flow-through was reapplied and centrifuged again. Columns were washed three times with buffer A and peptides were eluted with 3 x 2000 µl of 60% ACN into fresh tubes. The eluates were dried using an Eppendorf® Concentrator Plus (Eppendorf SE, Hamburg, Germany), and reconstituted in 2% ACN, 0.1% TFA. Peptide concentrations were determined using the Implen NanoPhotometer® N60/N50 (Implen GmbH, München, Germany). Samples were stored at -80°C.

### Phosphatase treatment heavy samples

For the phosphatase reaction heavy-labelled yeast peptide lysate was first mixed with phosphatase reaction buffer, then divided equally between two tubes. For peptide dephosphorylation with calf intestinal alkaline phosphatase (CIP), CIP was added to one aliquot (1 U CIP/µg peptide) while the other aliquot was left untreated. The reaction was carried out by incubating both CIP- and untreated samples at 37°C for 2 h. Following incubation, samples were quenched by adding TFA to a final concentration of 0.5% (v/v).

The samples were then desalted using STAGE-Tips^27^ as in the protocol described in our last publication.^16^

### Combining light & heavy CIP to generate mixed samples

After desalting, light and CIP-treated heavy peptide lysates were reconstituted in 2% ACN, 0.1% TFA. Peptide concentrations were determined using the Implen NanoPhotometer® N60/N50. Both lysates were combined in a ratio of 1:1 (m:m) to yield a mixed sample and divided equally between two tubes and dried again using the Eppendorf® Concentrator Plus at 45°C.

### Phosphatase treatment mixed samples

Phosphatase reaction was performed on the mixed samples as described above. CIP- and untreated mixed samples were desalted using STAGE-Tips and dried using an Eppendorf® Concentrator Plus.

All samples were reconstituted in 2% ACN, 0.1% TFA in preparation for LC-MS analysis. Peptide concentrations were determined using the Implen NanoPhotometer® N60/N50.

### LC-MS/MS analysis

LC-MS was performed using an Evosep One system coupled to a Tims TOF Ultra mass spectrometer equipped with a Captive Spray II ion source. 50 ng per sample were loaded on EvoTips according to the manufacturer’s recommendation. Peptide separation was conducted on an 8 cm × 150 µm ID Performance Column with 1.5 µm particle size, maintained at 40°C. A standard 60 SPD method was applied, utilizing a 21-min gradient. Solvent A consisted of 0.1% FA, and Solvent B was 80% ACN with 0.1% FA (Optima LCMS grade, Thermo Fisher Scientific, Waltham, MA, USA).

Data were acquired using a 2-frame Slice-PASEF scheme^15^ with a DIA cycle time of 0.89 s. The isolation window scheme spanned m/z 400−1200 and 1/K_0_ 0.75−1.3, covering most charge 2 precursor ions. The acquisition utilised 120 ms accumulation and ramp times. The window scheme was selected to balance sensitivity and coverage and consisted of one MS1 ramp and six MS/MS ramps, comprising a total of 90 MS/MS windows. The MS was operated in “high sensitivity detection mode”, optimised for low sample amounts.

### Raw data analysis

DIA-NN 2.2.0 was used to process the experiment, the full settings are reflected by the logs deposited to the PRIDE repository. Briefly, scoring was set to proteoforms. Mass accuracies were set to 15 ppm. The precursor’s m/z range was set to 300 to 1800 with a precursor charge range from 1 to 4. Samples were analysed using the following “additional options”: “--fixed-mod SILAC,0.0,K,label --lib-fixed-mod SILAC --channels SILAC,L,K,0; SILAC,H,K,8.014199 --peak-translation --original-mods --channel-run-norm.

For phosphopeptide identification the CIP-based spectral library described in our last publication was used^16^, allowing for up to three phosphorylation sites per peptide on S/T/Y. Scoring was set to proteoforms. Again, mass accuracies were fixed to 15 ppm. The precursor charge range was set to 1 to 4 and the precursor m/z range to 300 to 1800. As the same samples were processed in both studies, the library could be directly applied, substantially accelerating the search while maintaining high identification quality.

DIA-NN outputs were further filtered at Global.Q.Value < 0.01 and Channel.Q.Value < 0.01, unless indicated otherwise. For log_2_ standard deviation distributions, we optionally applied Quantity.Quality > 0.9.

Stoichiometry estimates for peptides quantified in both CIP- and untreated conditions were calculated using Precursor.Normalised quantities with the following transformation:

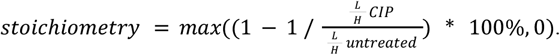

This metric reflects the pre-treatment phosphorylated fraction of each phosphopeptide. Across replicates, the median was calculated on log_2_-transformed L/H ratios.

For correlating SILAC-Site stoichiometries and phosphorylation stoichiometries estimated using ionisation efficiencies (IE) from Pho-Tip analysis,^16^ no Channel.Q.Value filtering was applied and Ms1.Apex.Area values were used. Log-scale total to unmodified ratios were calculated for untreated samples (light and heavy) analysed with the Pho-Tip library (observed phosphopeptides) using the formula:

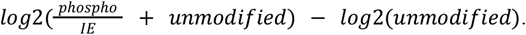

### Terminology and colour scheme

We established key peptide categories consistently applied throughout the manuscript, using a standardized colour scheme (Figure S1): “observed phosphopeptides” when searching SILAC-Site data with the Pho-Tip spectral library, “observed phosphosequences” when filtering the predicted library output for those phosphopeptide sequences, and “Pho-Tip phosphosequences” when filtering the predicted library output for all phosphopeptide sequences detected in the Pho-Tip dataset.

## Data availability

The MS proteomics data have been deposited to the ProteomeXchange Consortium via the PRIDE^28^ partner repository with the dataset identifier PXD078822.

## Author contributions

K.D.F., K.L.: experiments; K.D.F., V.D.: data analysis; L.R.S., V.D.: supervision; L.S.: mass spectrometry; K.D.F., V.D.: conception and results interpretation; K.D.F., L.R.S., V.D.: experiment design; K.D.F., L.R.S., J.R., V.D.: manuscript draft. All authors contributed to writing and approved the manuscript.

## Acknowledgements

This work is funded by the German Ministry of Education and Research (BMBF), as part of the National Research Node “Mass spectrometry in Systems Medicine” (MSCoreSys), under grant agreement 161L0221.

## Supplement

**Figure S1.**
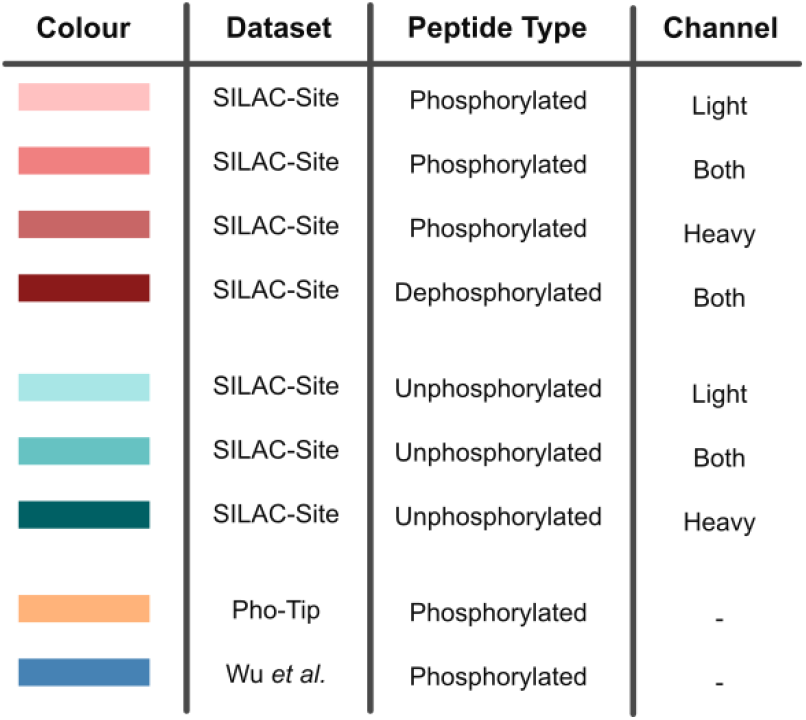
Colour Code. Colour palette used for all figures depending on dataset, peptide type and channel.

**Figure S2.**
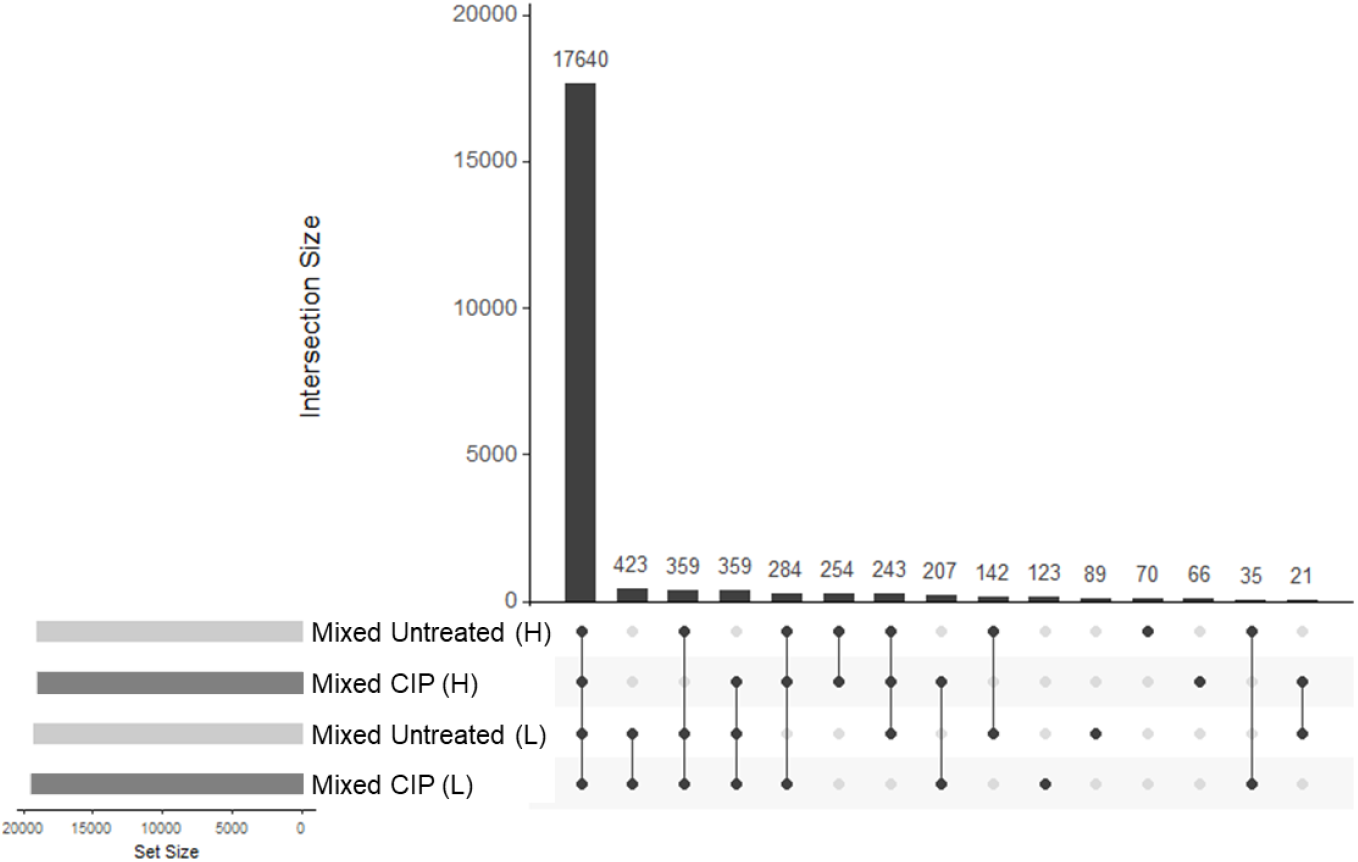
Consistent peptide identification across samples and channels. a) Uniquely identified peptide sequence intersections in mixed samples, stratified by treatment and channel.

**Figure S3.**
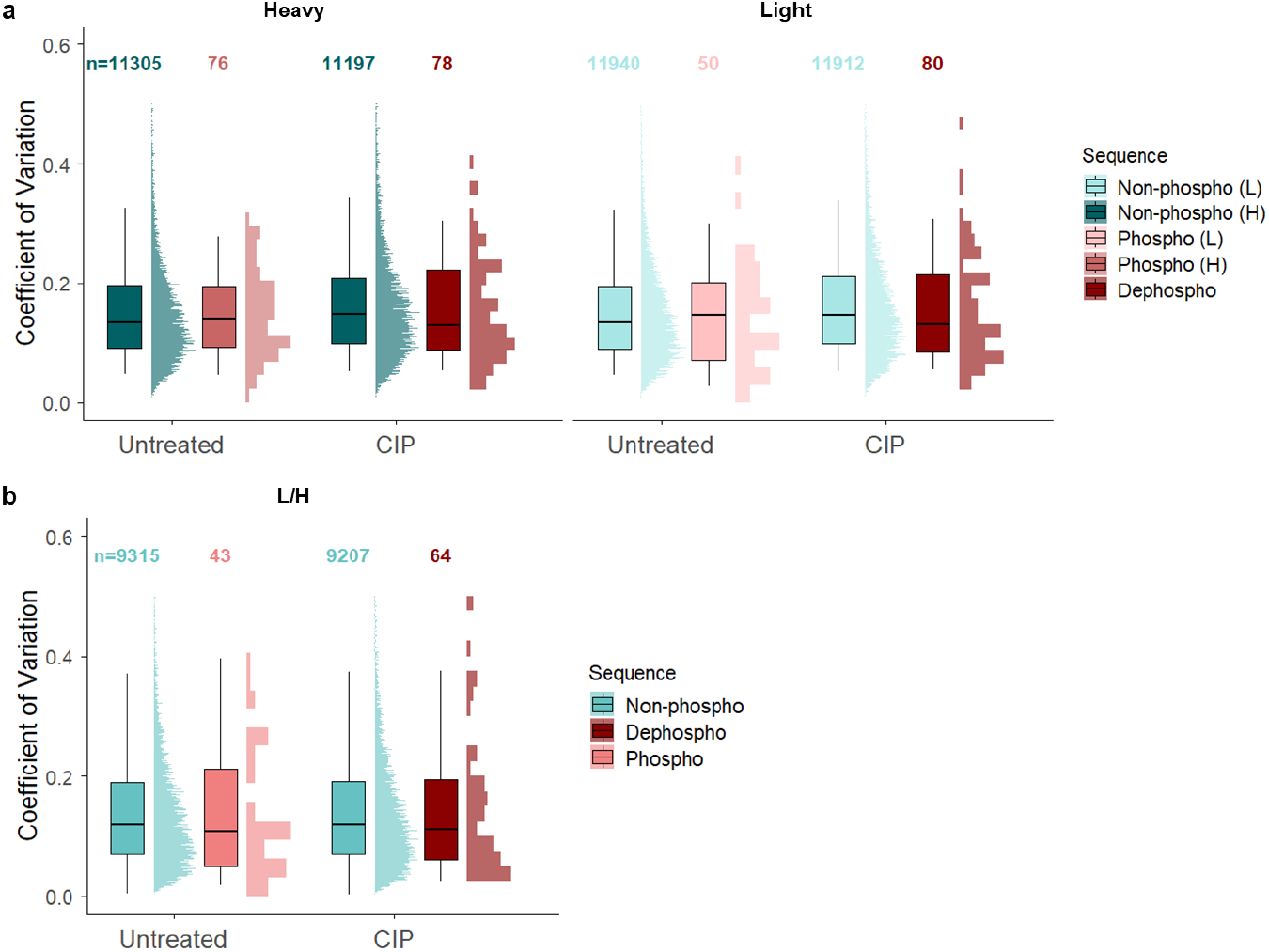
SILAC-Site quantitative reproducibility. a) Coefficient of variation (CV) values of peptide intensities across replicates, for peptide sequences identified in all replicates (n = 5), stratified by channel, treatment and phosphorylation status. b) Same analysis for CV values of L/H ratios across replicates. The boxes show the inter-quartile range (IQR) and the whiskers expand 1.5x the IQR.

**Figure S4.**
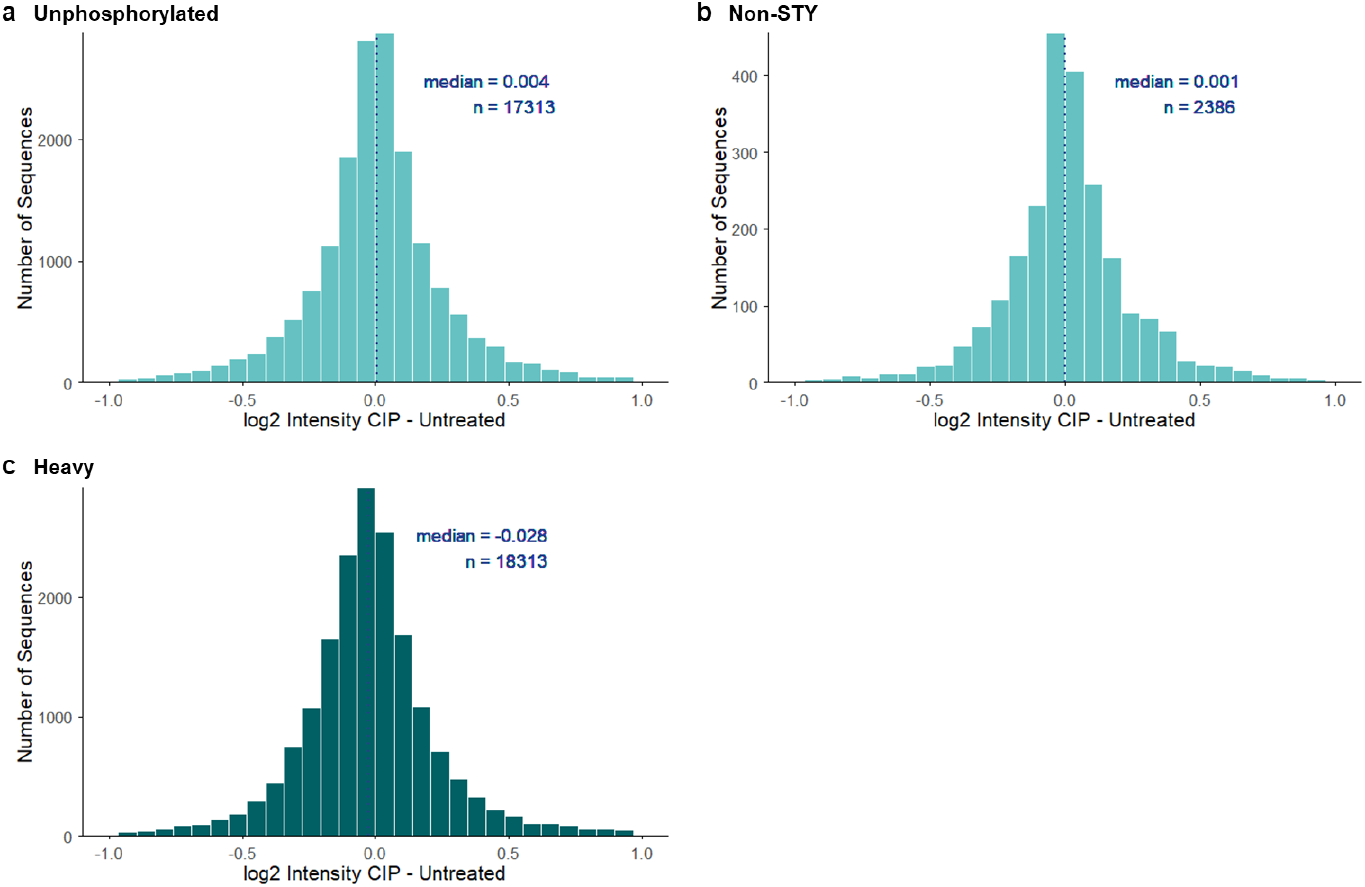
CIP treatment does not bias unphosphorylated peptide quantities. a) Distribution of log-scale L/H intensity shifts of unphosphorylated sequences detected in CIP- and untreated samples. b) Same analysis of non-phosphorylatable peptides lacking Ser/Thr/Tyr (STY) residues. c) Distribution of log-scale intensity shifts of all heavy channel unphosphorylated sequences detected in CIP- and untreated samples. Median values and number of peptide sequences are indicated for each distribution.

**Figure S5.**
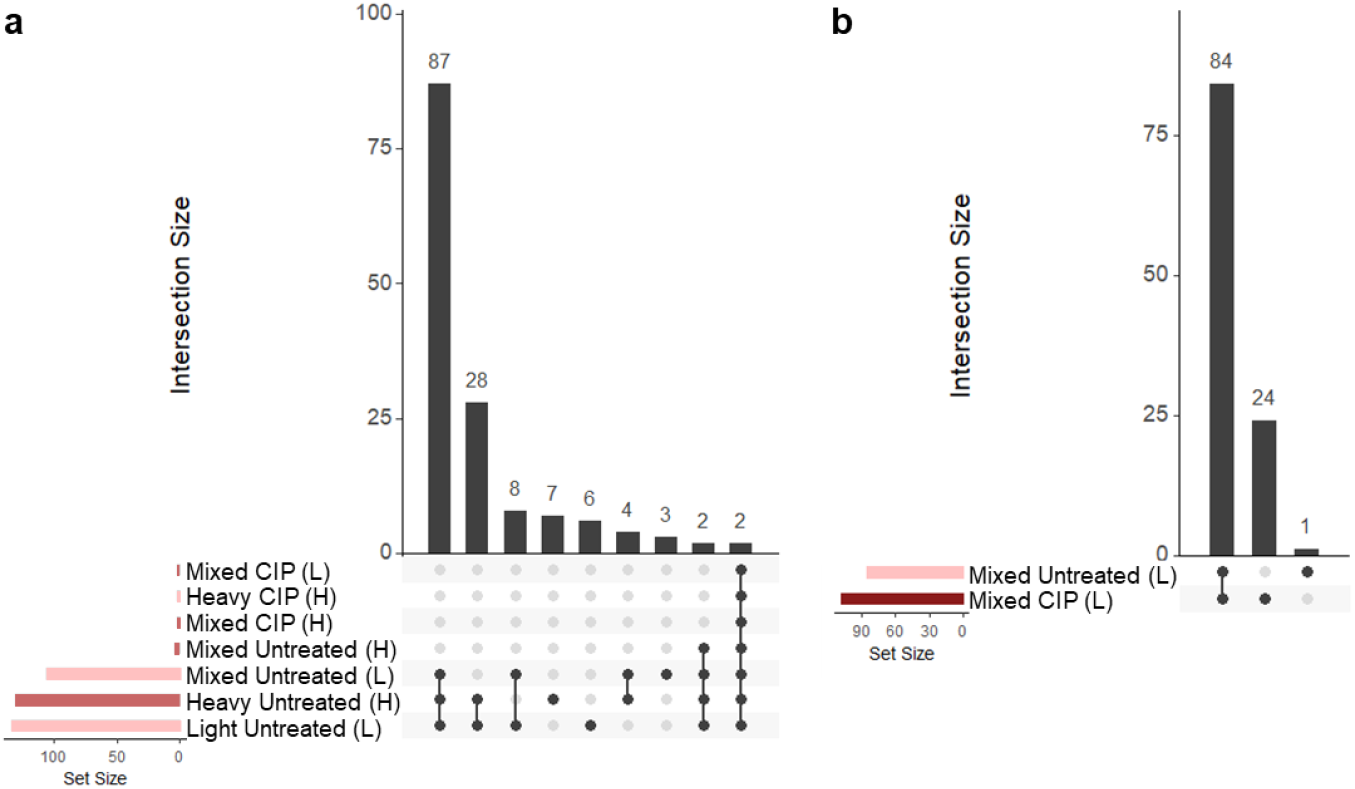
CIP treatment efficiently desphosphorylates peptides. a) Uniquely identified, observed phosphopeptides in untreated and their dephosphorylated counterparts in CIP-treated samples stratified by channel. b) Uniquely identified, observed phosphosequences detected in the untreated mixed sample (light channel), and their corresponding dephosphorylated counterparts in the CIP-treated mixed sample.

**Figure S6.**
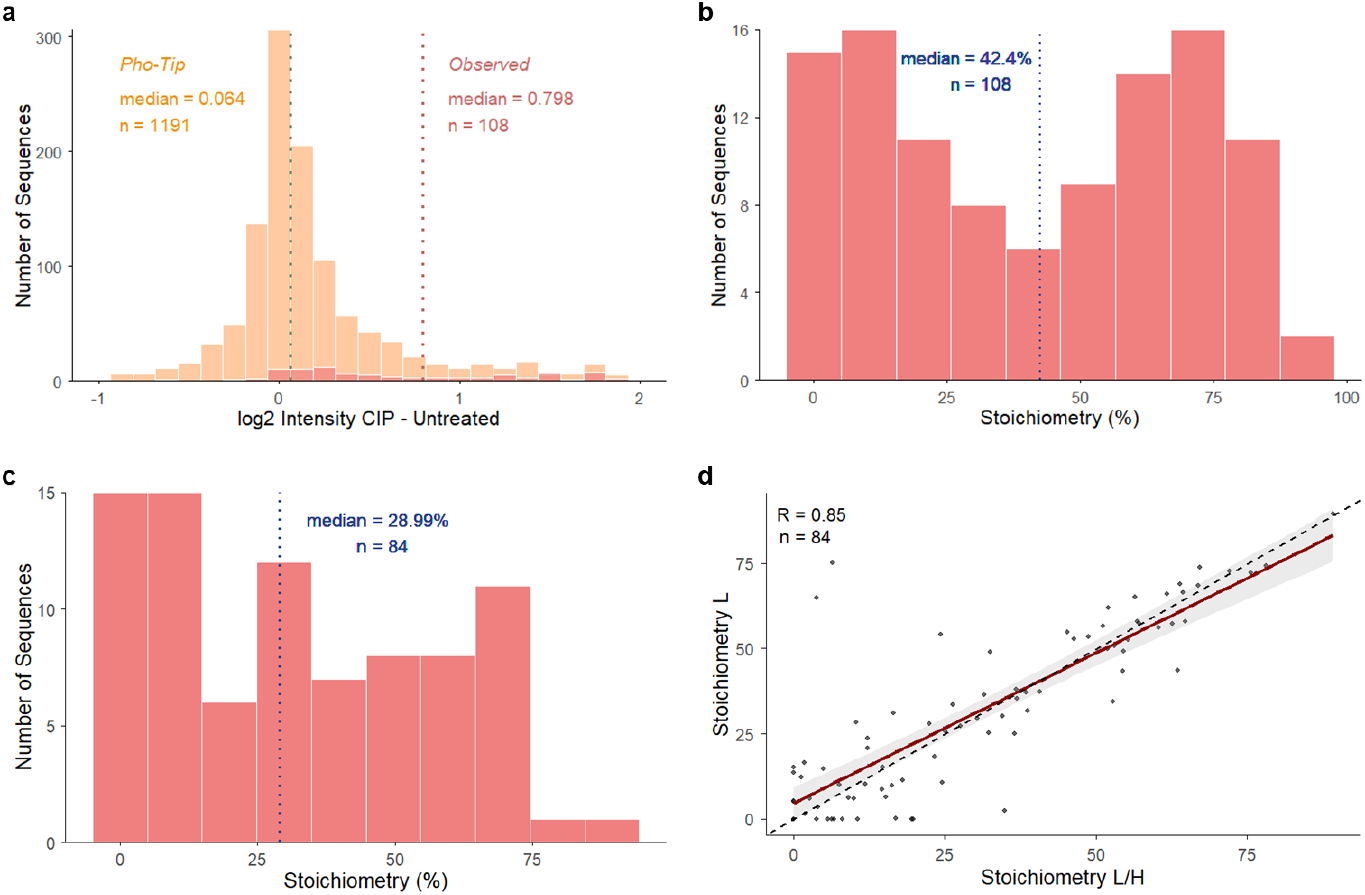
The effects of imputation and using light channel-only data on inferred stoichiometries. a) Distribution of log-scale (L/H) intensity shifts upon CIP treatment for observed and Pho-Tip phosphosequences detected in untreated and their dephosphorylated counterparts in CIP-treated samples. Missing values were imputed in the light channel in both, CIP- and untreated condition, with half minimum values. b) Phosphorylation stoichiometry (Methods) distribution inferred for observed phosphosequences using half-minimum value imputation. c) Phosphorylation stoichiometry (Methods) distribution inferred for observed phosphosequences, with stoichiometry estimates calculated using L channel intensities only. Median values and numbers of peptide sequences are indicated for each distribution (b, c). d) Correlation of observed phosphopeptides stoichiometries estimated from L/H ratios and L channel intensities only. Pearson correlation coefficient and number of peptide sequences are shown.

**Figure S7.**
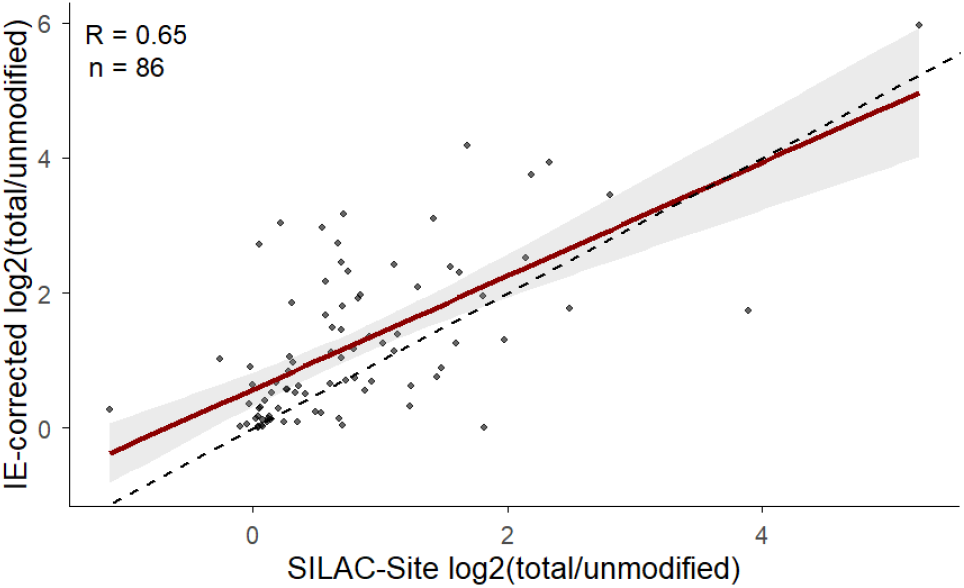
Orthogonal validation of stoichiometries inferred using SILAC-Site. Correlation of log_2_-transformed total to unmodified peptide amount ratios, estimated either by SILAC-Site (x-axis) or from measured MS1 signals for phosphorylated and corresponding unmodified peptides (y-axis) adjusted for their relative ionisation efficiencies derived from Pho-Tip analysis.^16^ Only peptides corresponding to observed phosphosequences were included. Pearson correlation coefficient and number of peptide sequences are shown.

**Figure S8.**
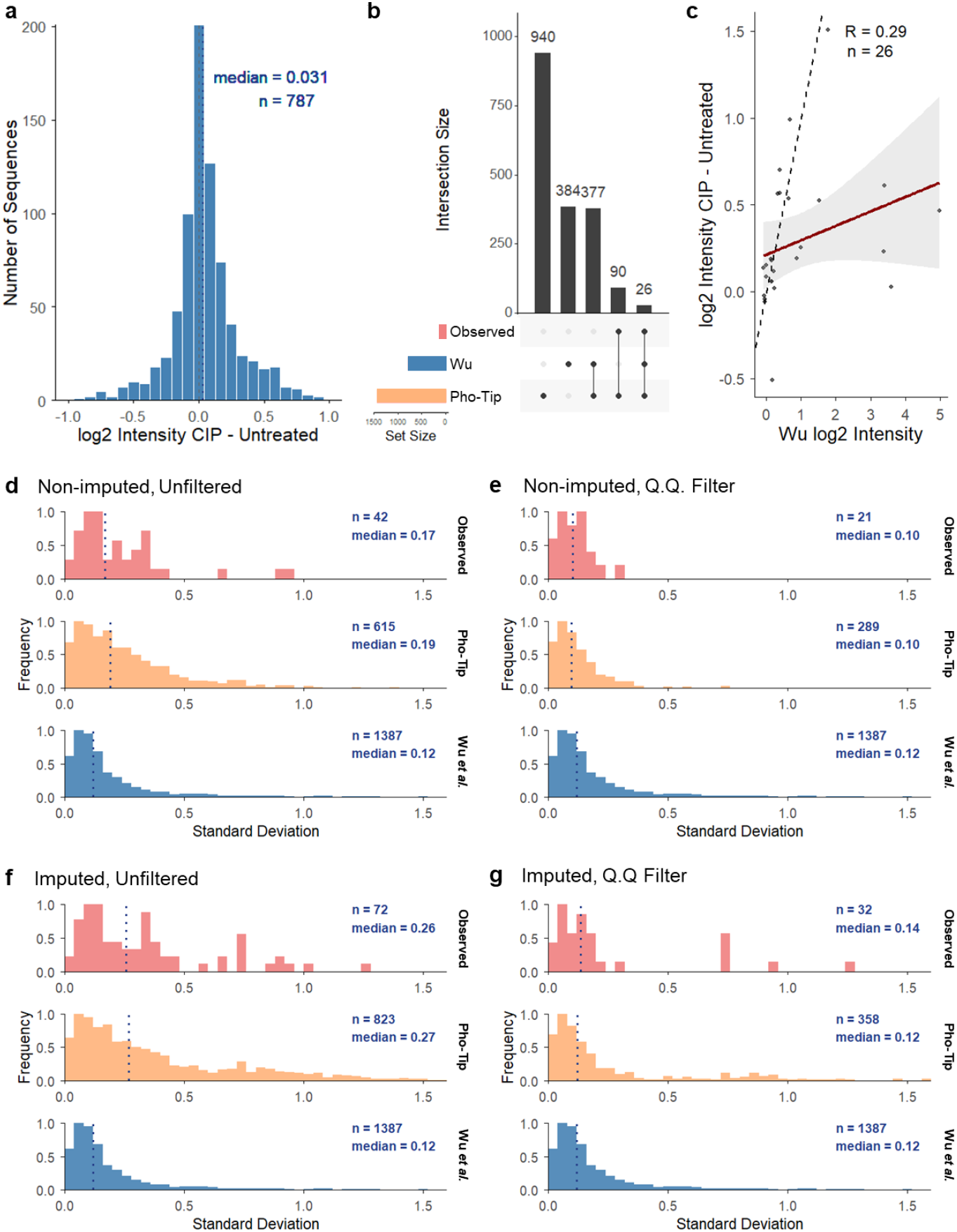
Comparison with Wu *et al* reference dataset. a) Distribution of log-scale (L/H) intensity shifts upon CIP treatment of all SILAC-Site peptide sequences filtered for phosphosites detected by Wu *et al*.^*4*^ b) Overlap of observed and Pho-Tip phosphosequences as well as all phosphosequences identified by Wu *et al*.^*4*^ c) Comparison of log_2_ intensity shifts upon dephosphorylation for observed phosphosequences and phosphosequences measured by Wu *et al*.^*4*^ d) Log_2_-space standard deviation distribution for phosphosequences detected in 3 experiments/replicates, separate for observed phosphosequences (red), Pho-Tip phosphosequences (orange) and phosphosequences measured by Wu *et al*^*4*^ (blue). e) Same analysis with Quantity.Quality > 0.9 filter (Methods) for SILAC-Site phosphosequences. f) Same analysis of imputed data without Quantity.Quality filter for SILAC-Site phosphosequences. g) Same analysis of imputed data with Quantity.Quality > 0.9 filter for SILAC-Site phosphosequences. Median values, Pearson correlation coefficient and numbers of peptide sequences are shown.

